# Isoniazid resistance in *Mycobacterium tuberculosis* is a heterogeneous phenotype comprised of overlapping MIC distributions with different underlying resistance mechanisms

**DOI:** 10.1101/524157

**Authors:** Arash Ghodousi, Elisa Tagliani, Eranga Karunaratne, Stefan Niemann, Jennifer Perera, Claudio U. Köser, Daniela Maria Cirillo

## Abstract

MIC testing using the BACTEC 960 MGIT system of 70 phylogenetically diverse, isoniazid-resistant clinical strains of *Mycobacterium tuberculosis* revealed a complex pattern of overlapping MIC distributions. Whole-genome sequencing could explain most of the level of resistance observed. The MIC distribution of strains with only *inhA* promoter mutations was split by the current concentration that is endorsed by the Clinical Laboratory Standards Institute to detect low-level resistance to isoniazid and is therefore likely not optimally set.

## Manuscript

In light of the continued selection and spread of drug-resistant tuberculosis, coupled with the dearth of novel antibiotics, the question of whether low-level resistance can be overcome by increasing the dose of a drug has become more urgent than ever (1). In 2018, the World Health Organization (WHO) has formally endorsed this possibility for moxifloxacin, whereby a dose of 800 mg/day can be used to treat low-level resistance to this fluoroquinolone, although the corresponding clinical breakpoint has not been recognized by the Clinical Laboratory Standards Institute (CLSI) (2-4). Conversely, for at least 15 years the position of CLSI has been to stratify resistance to the core drug isoniazid (INH) into low- and high-level resistance by testing two concentrations of this drug, whereas WHO has not endorsed this concept to date (5-7). Specially, the CLSI recommendation is to include the following statement in the antimicrobial susceptibility testing (AST) reports of strains that are only low-level resistant (i.e. that are resistant at the critical concentration but not the higher breakpoint of INH): “A specialist in the treatment of [multidrug resistant tuberculosis] should be consulted concerning the appropriate therapeutic regimen and dosages” (3). However, WHO is in the process of reviewing its recommendation for INH and, in its most recent manual for AST, has begun to stratify INH resistance on the genotypic level but has not yet set corresponding breakpoints to align the phenotype (6). We therefore set out to compare the phenotypic definitions of low- and high-level resistance by CLSI with the genotypic stratification proposed by WHO.

To this end, we used the BC BACTEC Mycobacteria Growth Indicator Tube (MGIT) 960 system to conduct comprehensive MIC testing of a selected set of phylogenetically diverse strains (70 INH resistant and 5 INH susceptible isolates, respectively) along with *M*. *tuberculosis* H37Rv ATCC 27294 as control strain. Four serial two fold *dilutions* were prepared from INH stock solution to provide a final test *range* of 0.016-0.25 μg/ml for susceptible and H37Rv strains, 0.25-4 μg/ml for *inhA* promoter mutant isolates with or without a concurrent *inhA* coding mutation, 1-16 μg/ml for S315T/N mutant isolates and 4-64 μg/ml for isolates with double mutations in *katG* and *inhA* promoter. The MGIT MIC was defined as the lowest antibiotic concentration that completely inhibited the growth of MTB when the control tube reached 400 growth unit. Whole Genome Sequencing (WGS) was performed using Nextera-XT DNA kit to prepare paired-end-libraries and sequenced on Illumina platform. Data analysis and SNPs calling were performed by MTBseq Pipeline (8) (for additional details see supplementary materials).

The six susceptible controls had INH MICs of 0.03-0.06 μg/ml (Figure 1 and Table S2). By contrast, resistant strains displayed a series of overlapping MIC distributions. Strains that only had mutations that are interrogated by the WHO-endorsed genotypic AST assays (i.e. the Hain GenoType MTBDR*plus* version 2 and Nipro NTM+MDRTB version 2) resulted in the following three MIC distributions (9). Strains that only had an *inhA* promoter changes or a mutation at codon 315 of *katG* had non-overlapping MIC distributions of 0.25-2 μg/ml and 4-16 μg/ml, respectively. Strains with both mutations displayed MICs of 8-64 μg/ml. The variation in these distributions was likely largely due to the normal variation in MIC testing (i.e. even in the same laboratory, a variation of plus or minus one dilution is inevitable, which is further exacerbated by the variation in testing between laboratories).

**Figure 1.**
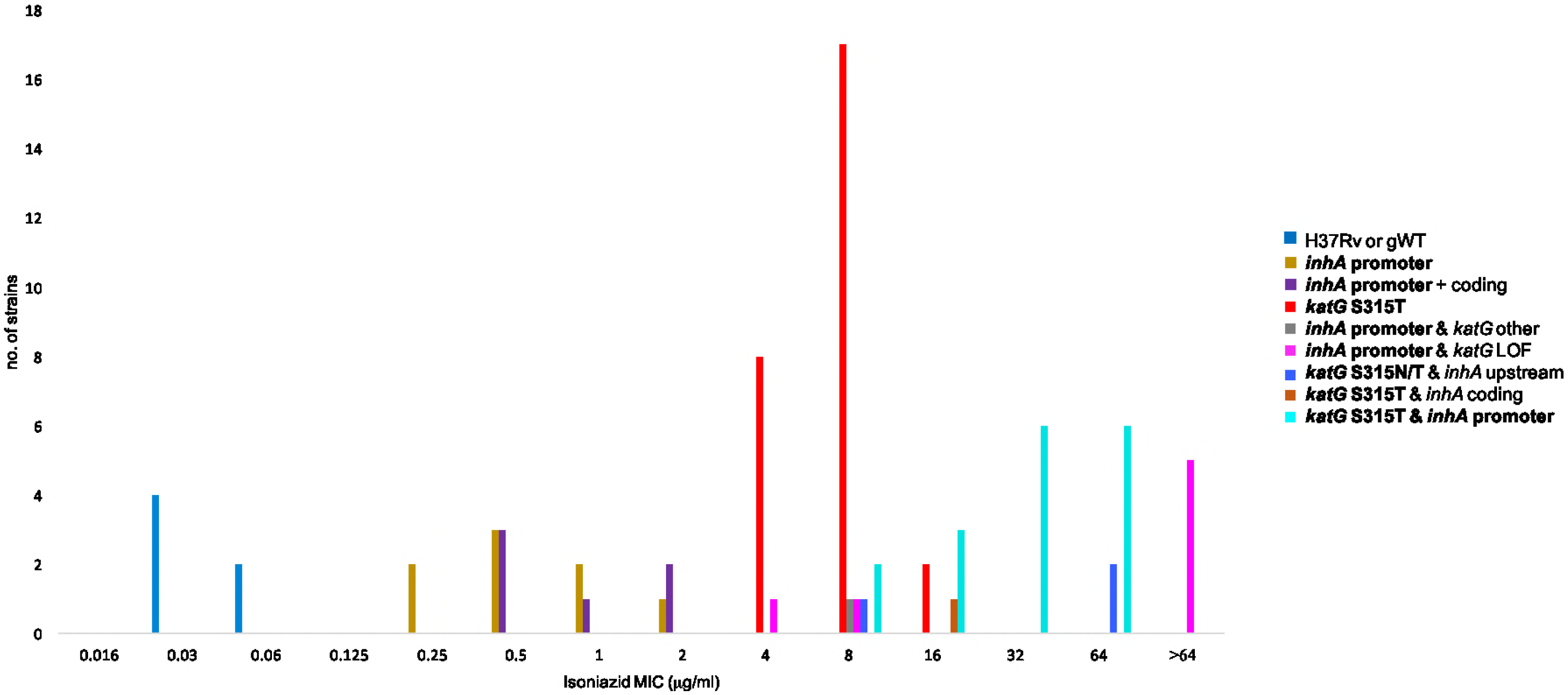
Isoniazid MIC results stratified by known or likely resistance mutations in the coding region of *katG* or *inhA*, or mutations that result in the overexpression of *inhA*. All of the latter mutations are upstream of *inhA*, but “promoter” was used to highlight the subgroup of these mutations (i.e. those in the -16 to -8 region upstream of the transcriptional start site of the *fabG1*-*inhA* operon (24)) that can be detected by the WHO-endorsed Hain GenoType MTBDR*plus* version 2 and Nipro NTM+MDRTB version 2 assays (all mutations interrogated by these assays are shown in bold in the legend of the plot).gWT=genotypically wild-type strains (i.e. strains without known resistance mutations); LOF=loss-of-function mutation (i.e. insertion, deletion, or nonsense mutation).

The precise level of resistance cannot be predicted using the Hain and Nipro assays alone because mutations that are not interrogated by these assays can increase the MICs. To some extent, this could be overcome by using WGS data, provided that know mutation or mutations with predictable effects were identified (i.e. the level of resistance could not be fully explained even with WGS). For example, some, but not all, strains with loss-of-function mutations in *katG* had MICs of >64 μg/ml (Figure 1). Moreover, a C deletion 34 nucleotides upstream of the main transcriptional start site of *inhA* likely accounted for the unusually high MIC of 64 μg/ml for a *katG* S315N mutant (10). By contrast, another mutation upstream of *inhA* at codon 203 of *fabG1*, which is known to result in the over-expression of *inhA* by creating an alternative promoter, did not appear to increase the MIC above the level explained by the *katG* S315T mutation in the strain in question (i.e. 8 μg/ml) (11). Similarly, there was an almost complete overlap between the MIC distributions of strains that harbor only *inhA* promoter mutations and those that had an additional *inhA* coding mutation at codons 21, 94, or 194 (i.e. 0.25-2 μg/ml compared with 0.5-2 μg/ml).

MGIT MIC data from more laboratories are needed to define robust quality control ranges/targets for INH and to quantify the effect of lab-to-lab variation on the MIC distributions described in this study (12). Assuming that our results are generalizable, several conclusions can be drawn. To assess the treatment efficacy of the standard or elevated dose of INH in the presence of a particular resistance conferring mutation, the upper end of the MIC distribution of the specific mutation has to be considered (13-15). For strain with only *inhA* promoter mutations, this target concentration would be 1 or 2 μg/ml (i.e. at least 10 times higher than the current critical concentration of 0.1 μg/ml (3, 6)). Should pharmacokinetic/pharmacodynamic, drug penetration, and clinical outcome data confirm that this target is achievable, it should be adopted as the clinical breakpoint as opposed to the current CLSI concentration of 0.4 μg/ml, which corresponds to 0.5 μg/ml using our dilution series (16, 17). This would avoid splitting the MIC distribution of *inhA* promoter mutants and would therefore reduce or eliminate their misclassification as high-level resistant because of the technical variation in AST, as would be the case with the CLSI breakpoint.

One argument against setting a clinical breakpoint at 1 or 2 μg/ml might be that it would result in the misclassification of strains with both *inhA* promoter and coding mutations as low-level resistant, as stressed in previous consensus statement (6, 18). However, two aspects should be borne in mind. First, only 3% (95% CI, 1-6%) of strains with *inhA* promoter mutations in the -16 to -8 region that do not have *katG* mutations have additional *inhA* coding mutations based on recent WHO population-level surveillance data from seven countries (19). Therefore, in most settings misclassifications of double mutants would be rare compared with the increased ability to detect *inhA* promoter mutants with a higher clinical breakpoint. Second, the effect of these coding mutations on the INH MIC and thus clinical outcome is likely modest at worst, but more MIC data are needed for the mutations at different *inhA* codons (20-23). Nevertheless, it might be advisable for countries that conduct routine WGS to err on the side of caution by classifying these double mutants as high-level resistant until clinical data to the contrary are available (in practice, however, it would be challenging to conduct a sufficiently powered study to address this question as these mutations are so rare).

